# Identification of antigen-specific TCR sequences using a strategy based on biological and statistical enrichment in unselected subjects

**DOI:** 10.1101/2020.05.11.088286

**Authors:** Neal P. Smith, Bert Ruiter, Yamini V. Virkud, Wayne G. Shreffler

## Abstract

Recent advances in high-throughput T cell receptor (TCR) sequencing have allowed for new insights into the TCR repertoire. However, methods for capturing antigen-specific repertoires remain an area of development. Here, we describe a novel approach that utilizes both a biological and statistical enrichment to define putatively antigen-specific complementarity-determining region 3 (CDR3) repertoires in unrelated individuals. The biological enrichment entails fluorescence-activated cell sorting of *in vitro* antigen-activated memory CD4^+^ T cells followed by TCRβ sequencing. The resulting TCRβ sequences are then filtered by selecting those that are statistically enriched when compared to their frequency in the autologous resting T cell compartment. Applying this method to define putatively peanut protein-specific repertoires in 27 peanut-allergic individuals resulted in a library of 7345 unique CDR3 amino acid sequences that had similar characteristics to validated antigen-specific repertoires in terms of homology and diversity. In-depth analysis of these CDR3s revealed 36 public sequences that demonstrated high levels of convergent recombination. In a network analysis, the public CDR3s unveiled themselves as core sequences with more edges than their private counterparts. This method has the potential to be applied to a wide range of T cell mediated disorders, and to yield new biomarkers and biological insights.

## Introduction

T cells are defined by their antigen-specific T cell receptor (TCR), and the collection of all unique TCRs in a human, which is dynamic and comprises approximately 10^10^ unique TCRs at a given time, is known as their TCR repertoire [1]. Unique antigen-specific TCRs are dimeric proteins comprising of an α and β chain that are both generated through the process of genomic rearrangement of germline V, D and J genes concurrent with random nucleotide insertions and deletions in the VDJ junction. This junction, known as the CDR3 (complimentary determining region 3), most closely interacts with an epitope during antigen presentation and therefore is the primary focus of studies aiming to elucidate the mechanisms of TCR specificity [2].

To date, ex-vivo analysis of antigen-specific TCRs has largely used selection of T cells by peptide-MHC (pMHC)-multimer (e.g. tetramer) binding and fluorescence-activated cell sorting (FACS). Of the 75,474 listed antigen-specific TCRs in the VDJdb (vdjdb.cdr3.net), 71,520 (94%) were reported to be isolated via pMHC-multimer selection [3]. In addition, new methodologies aimed at defining features that confer antigen-specificity have been benchmarked on various viral pMHC-tetramer selected TCRs [2, 4]. The advantage of tetramer selection is the high degree of specificity it provides as it captures cells specific for a single well-defined epitope. However, it comes with a major drawback in the form of a limited scope of analysis. Most immune responses are polyantigenic and current epitope mapping information of most antigens is incomplete. Additionally, pMHC-tetramers can only be used with T cell donors that have genetically matched HLA alleles. Thus tetramer-selection with only known epitopes in specific genetic backgrounds will likely provide an incomplete understanding of T cell mediated immune responses.

Public TCRs defined on the basis of identical amino acid sequences across multiple individuals have been described in many contexts since the early 1990s. Their existence has been intriguing given the vast number of possible recombination events, which suggests the prevalence of public TCRs would be much lower than what has been observed [5, 6]. One major mechanism that shapes an antigen-specific public TCR repertoire is convergent recombination, where selective pressure, such as that seen in response to dominant antigenic epitopes across individuals, gives rise to the presence of multiple unique nucleotide sequences (due to genetic code degeneracy and homologous gene segments) producing the same ‘public’ amino acid CDR3 [7]. Defining the public repertoire for antigen-specific responses can help in the development of diagnostics and therapeutics in T cell driven disorders.

Peanut allergy is a rising public health concern, and currently affects more than 1% of the US population. Compared with other food allergies, peanut allergy is less frequently outgrown and more often presents with severe symptoms [8]. Reactions to allergens are mediated by activation of mast cells and basophils through the high-affinity IgE receptor, occurring when receptor-bound specific IgE is cross-linked by binding to peanut allergens. Production of this high affinity allergen-specific IgE is T cell-dependent, and a peanut-specific transcriptional profile characterized by increased expression of IL-4, IL-5, IL-9 and IL-13 has been observed in CD4^+^ T cell subsets from subjects with peanut allergy [9] [10]. Multiple forms of immunotherapy for the treatment of peanut allergy are currently in clinical trials, of which oral immunotherapy is the most common. The effects of these treatments on the peanut-specific CD4^+^ T cell response and their correlation with clinical success are an active area of investigation [11] [12].

In this work, we investigated the antigen-specific TCR repertoires in peanut-allergic individuals. We isolated peanut protein-activated and resting memory CD4^+^ T cells and sequenced the CDR3 of the TCR β-chain (CDR3β). The most enriched sequences in the peanut-activated compartment were identified as putatively peanut-specific CDR3βs (ps-CDR3s). These biologically and statistically selected sequences exhibited properties associated with antigen-specific populations, such as increased homology, decreased diversity and instances of convergent recombination. Within our pool of ps-CDR3s, we found a subset of sequences enriched in multiple subjects (i.e. public ps-CDR3s), suggesting the existence of public T cell responses among peanut-allergic individuals with diverse HLA genotypes. This work describes a novel method to study antigen-specific TCR repertoires when information on epitopes and/or HLA genotypes is not available or incomplete. This method is potentially applicable to studies of various T cell-mediated diseases beyond the scope of allergy.

## Results

### Selection of putatively peanut-specific CDR3βs

With the goal of enriching for antigen-specific memory CD4^+^ T cells, peripheral blood mononuclear cells (PBMCs) from 27 peanut-allergic individuals were cultured for 20 hours with a peanut protein extract. The activation marker CD154 (CD40L) was used to discriminate between peanut-activated CD154^+^ and resting CD154^−^CD69^−^ memory T cells, and TCR CDR3β sequences were generated from both populations (Fig. 1A, Supplemental fig 1) [13, 14]. Flow-cytometric analysis revealed that stimulation with peanut protein increased the expression of CD154 on CD4^+^ memory T cells as compared to unstimulated cultures (median[IQR] CD154^+^ cells per million CD4^+^ T cells: 2504[1328, 4105] stimulated vs. 172[117, 319] unstimulated; P < 0.0001) (Fig. 1B). To filter out sequences that were more likely to be present in the activated population due to bystander activation, we developed a statistical enrichment strategy to define putatively peanut-specific CDR3 sequences (ps-CDR3s). First, a *G*-test of independence was applied to all CDR3 sequences in the peanut-activated compartment and adjusted P-values (q-values) were generated based on a FDR of q<0.05. Sequences were further filtered to include only those with read counts ≥2 in the activated compartment and those whose proportion in the activated compartment was higher than that of the resting compartment. This approach defined 7345 out of the 53205 unique amino acid sequences in the peanut-activated compartment (14%) across our 27 peanut-allergic individuals as ps-CDR3s. While the vast majority (n = 7309) of ps-CDR3s were private, 36 unique amino acid sequences (0.49% of ps-CDR3s) were found in more than one individual (i.e., public). Of these public sequences, 33 were found in only two individuals. However, one ps-CDR3 (CASSFRFLSGRALNEQFF) was statistically enriched in 8 of the 27 subjects, suggesting a strong selective pressure for entrance of this sequence into the peanut-activated memory T cell repertoire (Supplemental fig 2). The CDR3β amino acid- and nucleotide-sequences that met our ps-CDR3 enrichment criteria only represented a minor fraction of the entire CDR3β pool from peanut-activated T cells (amino acid: median 14.05%, nucleotide: median 14.46%), indicating a high level of stringency with this statistical approach (Figure 1C). The median frequency of ps-CDR3s was 117 per million CD4^+^ T cells (IQR = 46.8, 173.15) which corresponds well to published frequencies of whole allergen-specific T cells in allergic individuals, suggesting the stringency is appropriate (Figure 1D) [9, 15, 16].

**Figure 1:**
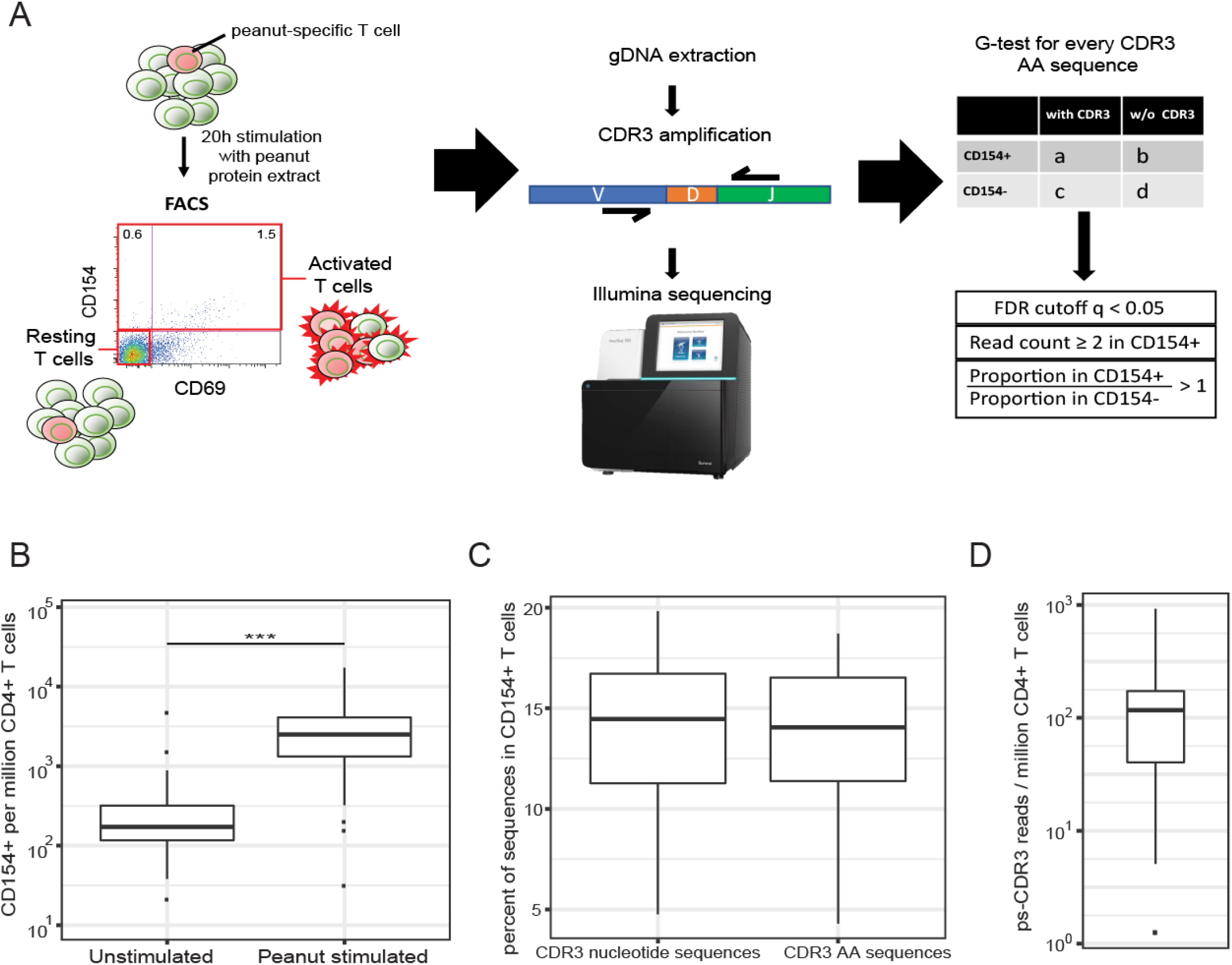
Selection of putatively peanut-specific CDR3s (ps-CDR3s). A, A general schema for the selection of ps-CDR3s (see methods). B, The frequency of CD154^+^ T cells per million CD4^+^ T cells in peanut protein-stimulated PBMC cultures is higher than in unstimulated cultures (n = 27, *** P < 0.001, Wilcoxon matched-pairs signed rank test). C, Percentage of CDR3s in the CD154^+^ T cell compartment that met the criteria to be selected as ps-CDR3s. Boxplots represent the percentage of unique CDR3 nucleotide (left) and amino acid (right) sequences that were ps-CDR3s (n = 27). D, The frequency of ps-CDR3s per million CD4^+^ T cells in peanut protein-stimulated cultures (n = 27).

### ps-CDR3s are more oligoclonal and similar than unselected CDR3s from both activated and resting T cells

To assess the efficacy of our statistical enrichment method for selecting those sequences most likely to be peanut-specific, we examined the similarity of the ps-CDR3s using Hamming and Levenshtein distances. For each ps-CDR3, the minimum Hamming and Levenshtein distances were measured by determining the minimum number of amino acid differences compared to its next closest ps-CDR3. As a control, the minimum Hamming and Levenshtein distances of equal-sized random repeated samplings of resting (CD154^−^) and activated (CD154^+^) CDR3s were also measured (figure 2A, B). There were significantly more sequences with a minimum Hamming distance of 0-2 and Levenshtein distance of 0-1 in the ps-CDR3s than in either the activated or resting CDR3βs (P < 0.01), indicating enhanced similarity among the statistically enriched sequences. Importantly, these metrics showed no differences between random activated vs. resting CDR3s, supporting our hypothesis that there are many non-peanut-specific sequences in the activated pool under this methodology and that our statistical enrichment strategy is effective in removing them.

**Figure 2:**
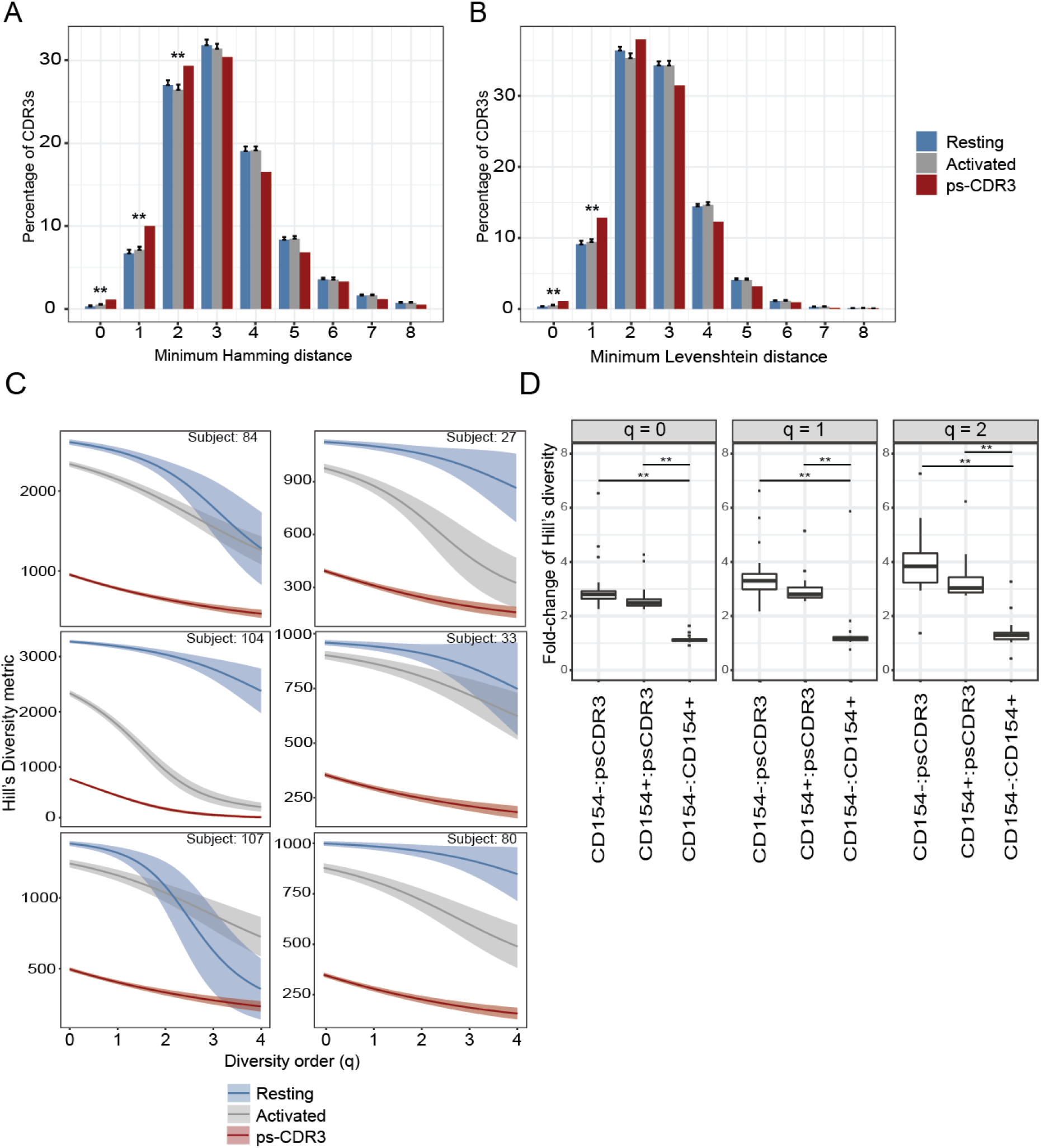
ps-CDR3s demonstrate increased similarity and decreased diversity when compared to total activated and resting CDR3s. A-B, Minimum Hamming- and Levenshtein-distance of ps-CDR3s compared to 100 equal-sized random-resamplings of total activated and resting CDR3s (** P < 0.01, Fisher’s exact test). C, Smoothed Hill’s diversity curve at diversity orders 0-4 for 6 subjects. Diversity of ps-CDR3s was significantly lower than that of activated and resting CDR3s at all diversity orders (P < 0.01, empirical cumulative distribution function of bootstrap delta distribution, see methods). D, Fold-change of Hill’s diversity metric of ps-CDR3:activated, ps-CDR3:resting, and activated:resting sequences. Distributions represent values from 25 subjects with > 30 unique ps-CDR3s (** P < 0.001, Wilcoxon matched-pairs signed rank test).

Uniform resampling of autologous sets of ps-CDR3s, activated and resting sequences were used to measure Hill’s diversity index across different diversity orders to capture both the true richness (low q value) and abundance (high q value) of the samples (figure 2C) [17]. For the 25 subjects with >30 ps-CDR3s, the general diversity index was significantly lower for the ps-CDR3s than for either the activated or resting sequences at diversity orders 0-2 (P < 0.01). Consistent with biological enrichment, the diversity of the activated sequences was significantly lower than that of the resting samples at diversity orders 0-2 for 22 of the 25 subjects (P < 0.01). Furthermore, the difference in diversity between the ps-CDR3s and either the activated or resting CDR3s was significantly greater than the difference observed between the activated and resting CDR3s (P < 0.001), emphasizing the utility in our statistical filtering (figure 2D).

### Public ps-CDR3s show evidence of convergent recombination

Antigen-specific public TCRs from individuals with different HLA genotypes can provide valuable insights into shared immune responses within a diverse population. For this reason, we wanted to further examine the characteristics of our public ps-CDR3s. We observed that the public ps-CDR3s contained fewer total N-nucleotide insertions than the private ps-CDR3s (median public: 3 insertions, median private : 7 insertions; P < 0.01). Similar patterns were observed when looking specifically at V-D insertions (median public : 1 insertion, median private : 3 insertions; P < 0.01) and D-J insertions (median public : 1 insertions, median private : 3 insertions; P < 0.01) which is consistent with prior literature demonstrating that public TCRs are closer to germline (figure 3A) [18, 19] [20] [7].

**Figure 3.**
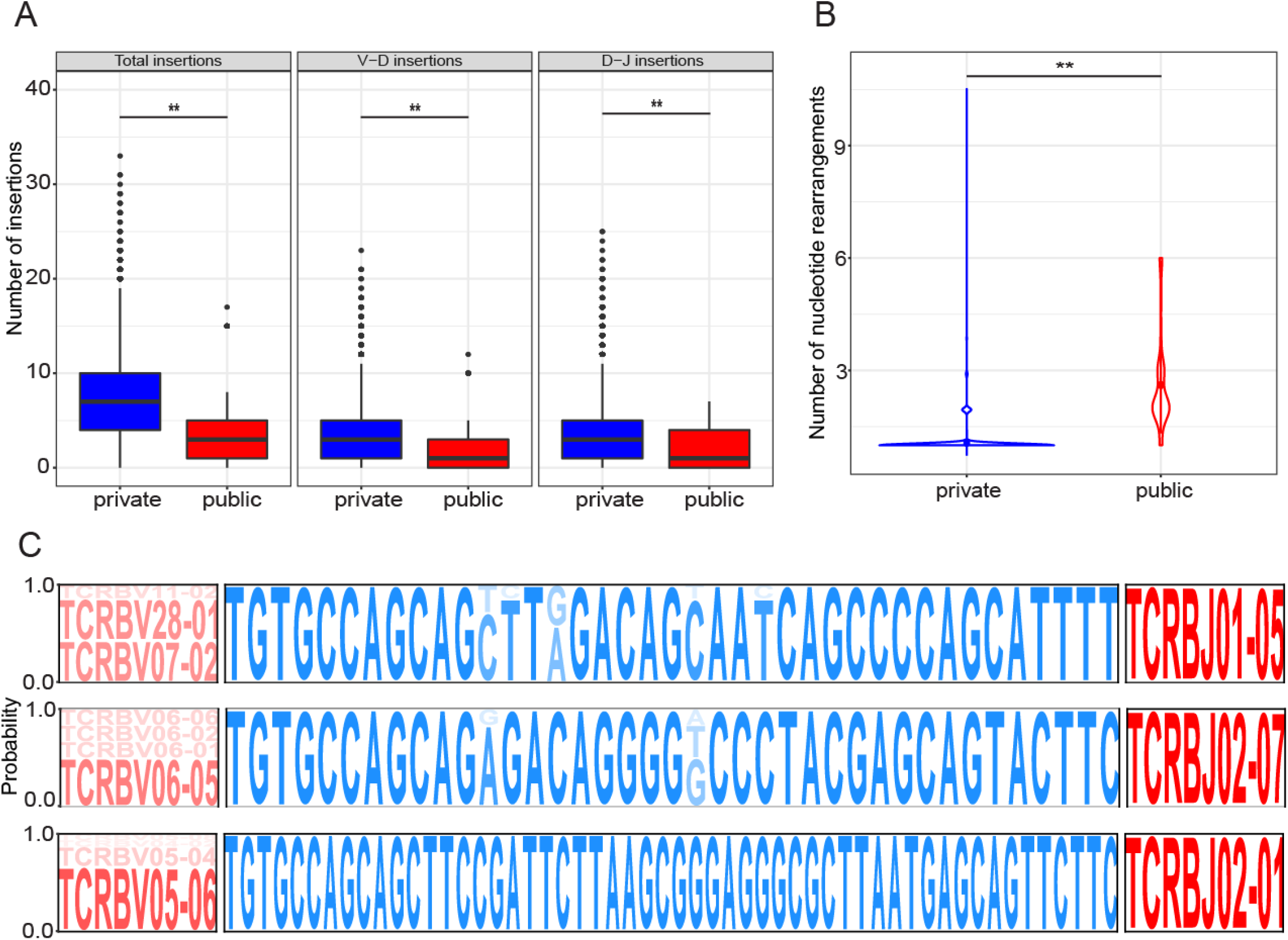
Public ps-CDR3s are characterized by being closer to germline and exhibiting convergent recombination. A, Public ps-CDR3s are closer to germline than private ps-CDR3s. Distribution of total, V-D and D-J insertions of public and private ps-CDR3s (P < 0.01, Mann-Whitney U test). B, Public ps-CDR3s demonstrate more convergent recombination than private ps-CDR3s. Violin plots represent the distribution of unique nucleotide rearrangements responsible for each unique public and private ps-CDR3 (P < 0.001, Mann-Whitney U test). C, Convergence of public ps-CDR3s occurs in both germline (V-gene) and non-germline (CDR3) regions. Logo plots represent the probability of either specific gene segments (red) or CDR3 nucleotides (blue) to be found for their corresponding amino acid sequence.

Convergent recombination of TCRs (multiple TCR nucleotide sequences translating to the same amino acid sequence) provides evidence of antigenic selection [7]. We searched the ps-CDR3s for this phenomenon and observed that convergence occurred in 467 out of 7345 unique ps-CDR3 amino acid sequences (6.3%). Interestingly, convergence was very common among the public ps-CDR3s, occurring in 33 out of 36 public amino acid sequences (91.6%), which suggests a strong selective pressure for these sequences’ entrance into the peanut-activated memory T cell pool. Additionally, the number of nucleotide rearrangements in the public ps-CDR3s was significantly higher than that in private ps-CDR3s (P < 0.001) (figure 3B). Analysis of the most convergent public sequences (≥ 5 unique nucleotide rearrangements) demonstrated convergence derived from germline (V-gene) and non-germline (N-nucleotide insertions) changes (figure 3C).

### Analysis of ps-CDR3 amino acid sequences reveals public, dominant motifs

Given the ps-CDR3 sequences demonstrated enhanced overall similarity, a motif analysis was performed according to Glanville et al. [2] to find dominant patterns among the putatively peanut-specific pool of sequences (supplemental figure 3A). Briefly, for all ps-CDR3s, the region most likely to be in contact with an antigenic peptide was broken into continuous 3mer, 4mer and 5mer amino acid sequences. In addition, discontinuous 4mers and 5mers were generated to allow for the detection of motifs with gaps. Nmers that were at least 10-fold enriched in the ps-CDR3s as compared to the resting sequences and found in at least 3 unique ps-CDR3s from at least 3 individuals, were considered dominant motifs (figure 4A-B, supplemental figure 3B-D). These criteria were met by 148 of the 104520 unique nmer sequences (0.14%), and the proportion of dominant motifs in the total motif pool was fairly similar for each nmer size (0.06 – 0.16 %) (table 1). With these strict criteria, 399 of the 7345 ps-CDR3s (5.4%) contained dominant motifs, which likely represent sequences specific for the most immunogenic T cell epitopes in peanut protein. Indeed, there were significantly more public ps-CDR3s (16.6%) than private ps-CDR3s (5.3%) that contained a dominant motif, supporting the hypothesis that these motifs are associated with the most common epitopes (P < 0.05) (figure 4C).

**Table 1:**
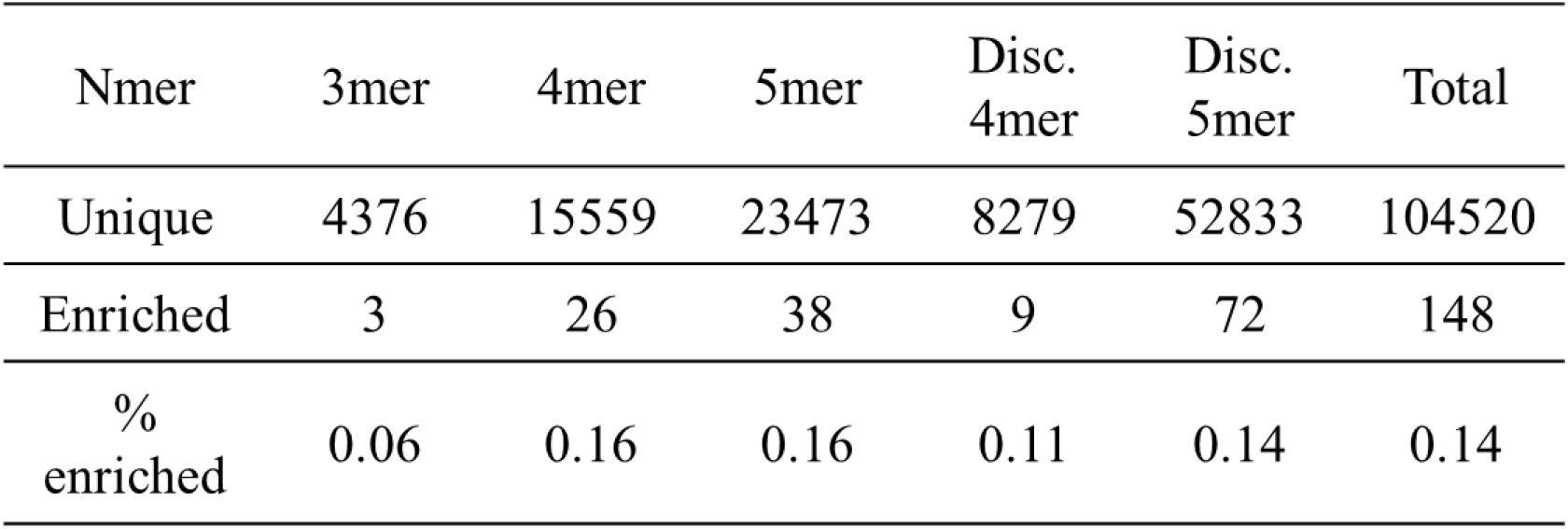
Summary of all nmers generated from the ps-CDR3s.

**Figure 4.**
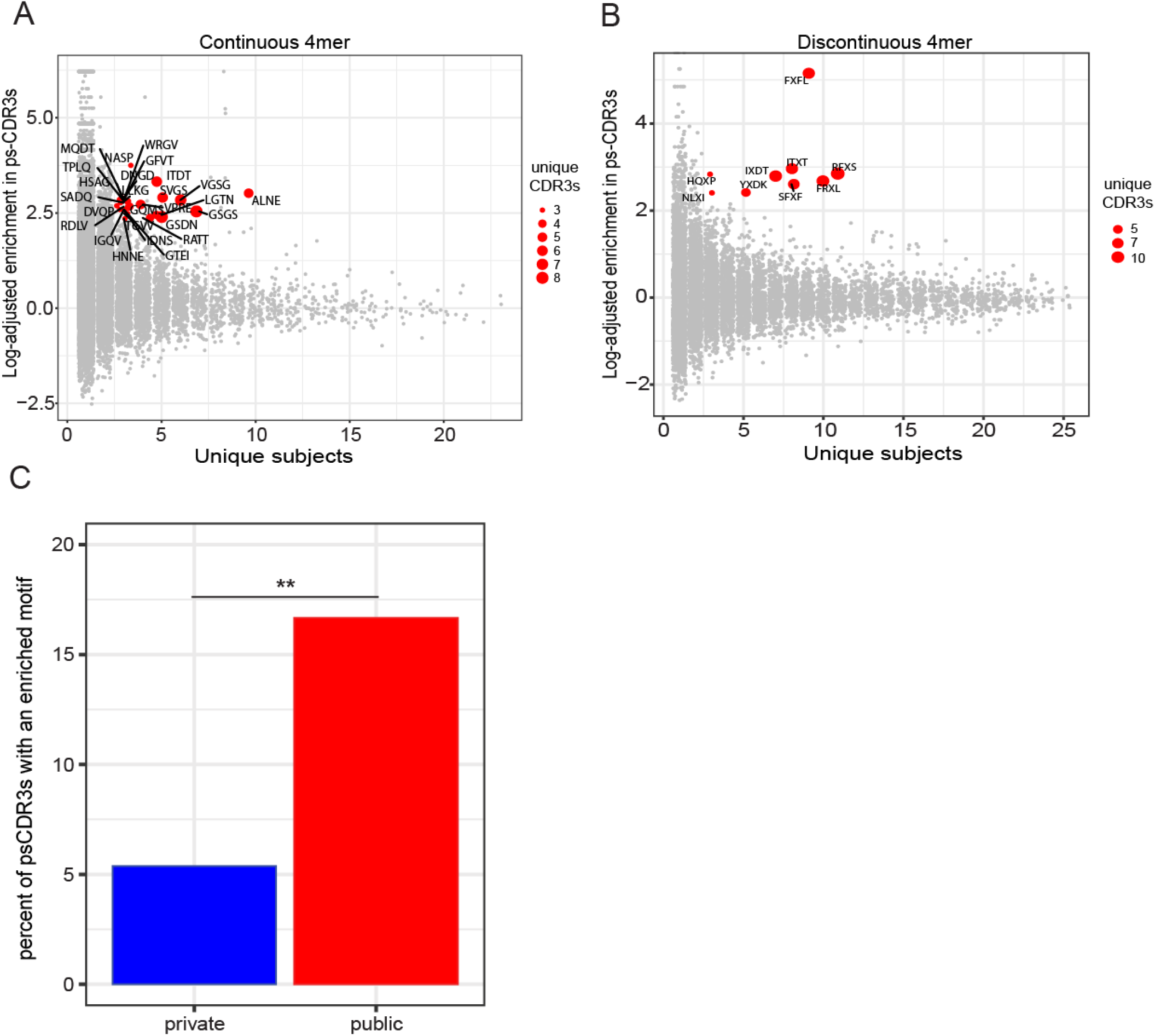
ps-CDR3s contain public, dominant motifs. A-B, The ps-CDR3s contain shared, enriched motifs as compared to resting CDR3s. Red points represent the most dominant motifs, found to be ≥10-fold enriched in ps-CDR3s:resting CDR3s, and present in ≥ 3 unique ps-CDR3s derived from ≥ 3 subjects. C, Public ps-CDR3s are more likely to have a dominant motif than private ps-CDR3s (P < 0.05, Fisher’s exact test).

### Network analysis confirms that public ps-CDR3s are core sequences

Utilizing the measured metrics of similarity, the dominant motifs and evidence of convergent recombination, we performed a network analysis to group highly homologous sequences (figure 5A). Similar to previously published approaches [2, 21, 22], each ps-CDR3 amino acid sequence was treated as a node and an edge was created between nodes if the ps-CDR3s were within a Levenshtein distance of 1 or shared a dominant motif. In addition, self-edges were created to represent additional nucleotide sequences for a corresponding amino acid sequence. Overall, there were 1759 edges created among 1324 ps-CDR3s. Interestingly, the majority (n = 23; 63.8%) of public ps-CDR3s were present in the network, whereas a much smaller fraction of private ps-CDR3s (n = 1301; 17.79%) were found in the graph (P < 0.001) (figure 5B). Moreover, the node degree (the number of edges a node has) of the public sequences was higher than that of the private sequences (public median node degree: 4, private median node degree: 2; P < 0.001) (figure 5C). To assess the amount of structure observed among the ps-CDR3s, we compared the number of edges in the ps-CDR3 graph to the median number of edges created when performing the same network analysis on equal-sized random repeated samplings of activated or resting CDR3s. Indeed, there were significantly more edges created between the ps-CDR3s than between activated CDR3s (median 1227 edges) or resting CDR3s (median 1107 edges) (P < 0.001) (figure 5D). We hypothesized that specific clusters of CDR3s would be associated with individual HLA genotypes, which are shown in supplemental table 1. However, no correlations could be found. Taken together, these data suggest that public ps-CDR3s represent “core” sequences that likely bind dominant epitopes recognized by multiple individuals who are peanut-allergic.

**Figure 5.**
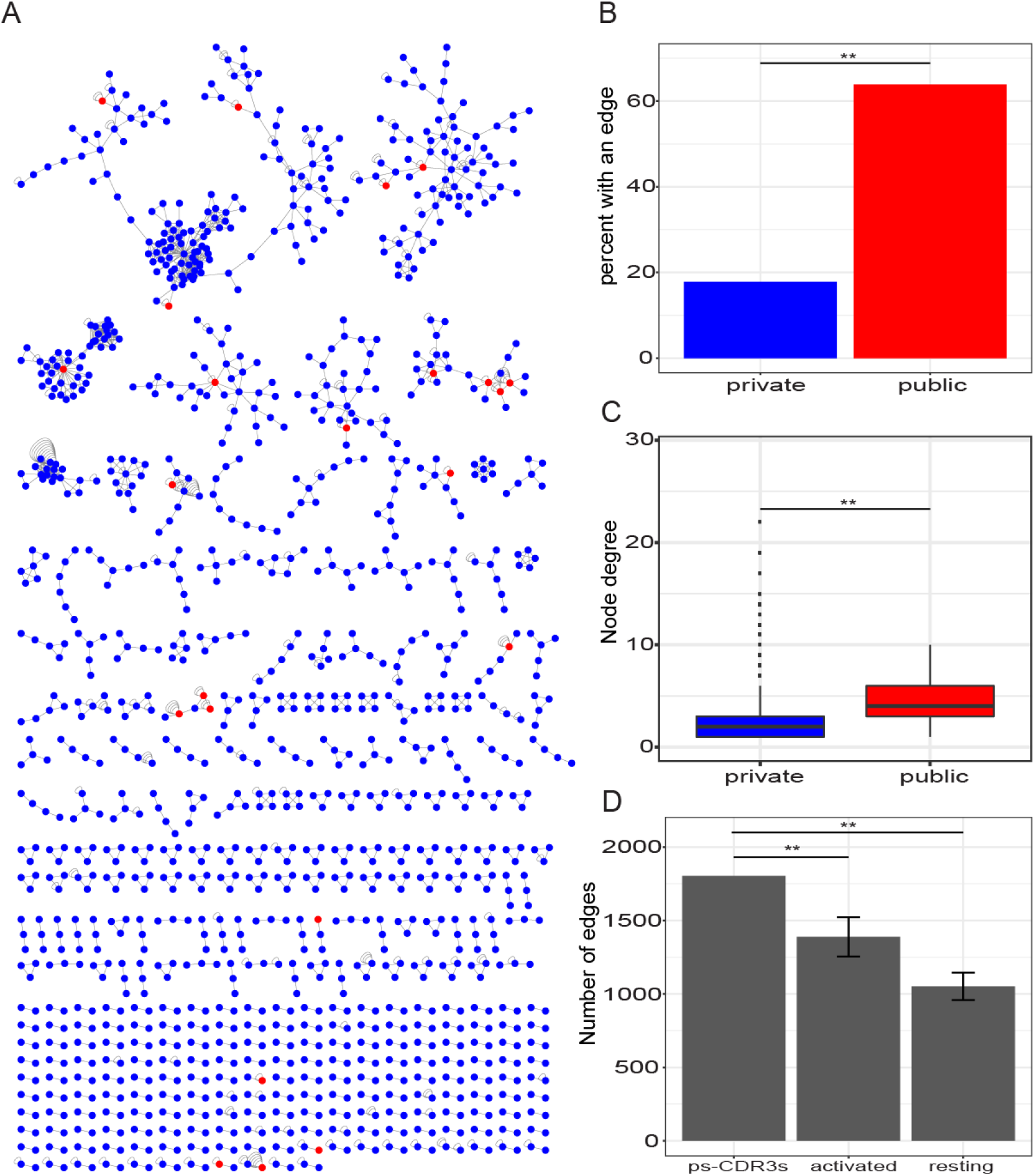
Network analysis reveals that public ps-CDR3s are core sequences with more structure than activated and resting CDR3s. A, Network analysis of ps-CDR3s where each node represents a unique ps-CDR3 amino acid sequence. Edges were created between the nodes if they were within a Levenshtein distance of 1 or if the nodes shared an enriched motif. Selfedges were created on nodes to represent additional unique nucleotide sequences for each unique amino acid sequence (i.e. convergence). Blue nodes represent private ps-CDR3s and red nodes represent public ps-CDR3s. B, The percent of public ps-CDR3s with an edge to another ps-CDR3 (as shown in panel A) was higher than that of private ps-CDR3s (**P < 0.001, Fishers exact test). C, The overall node degree of public ps-CDR3s was higher than that of private ps-CDR3s. Boxplots represent distribution of the node degree of all private (blue) and public (red) ps-CDR3s in the graph (** P < 0.001, Mann-whitney U test.). D, The number of total edges created among the ps-CDR3s compared to the median number of edges created among 100 equal-sized random resamplings of total activated and resting CDR3s. Error bars represent standard deviation. (** P < 0.001, z-test).

### ps-CDR3s are more likely to have a T_eff_ phenotype than a T_reg_ phenotype

To better understand the phenotype of the CD4^+^ T cells from which the ps-CDR3s were derived, TCRβ-sequencing was performed on bulk CD25^+^CD127^+^ effector T cells (T_eff_) and CD25^+^+CD127-regulatory T cells (T_reg_) from 9 of the 27 subjects used in the initial analysis (supplemental figure 4A). These bulk sequences were then probed for ps-CDR3 amino acid sequences. Although we obtained more T_reg_ sequences overall, ps-CDR3s were more commonly found in the T_eff_ compartment (figure 6A; Supplemental figure 4B-C), indicating either a bias in the induction of CD154 that favors the detection of peanut-specific T_eff_ over T_reg_, or a propensity of the peanut-specific T cell compartment of allergic individuals to be dominated by T_eff_. Interestingly, the imbalance between T_eff_ - and T_reg_-derived sequences was enhanced by our statistical enrichment, indicating that the most enriched sequences in the activated compartment are predominantly T_eff_ (P < 0.05) (figure 6B). The percentage of public ps-CDR3s found in the T_eff_ compartment was significantly higher than that of the private ps-CDR3s (public: 63.8%, private: 24.1%; P < 0.01). A similar result was found in the T_reg_ subsets (public: 36.1%, private: 10.72%; P < 0.01). These observations confirm that our public ps-CDR3s represent sequences common among peanut-allergic individuals (figure 6C).

**Figure 6.**
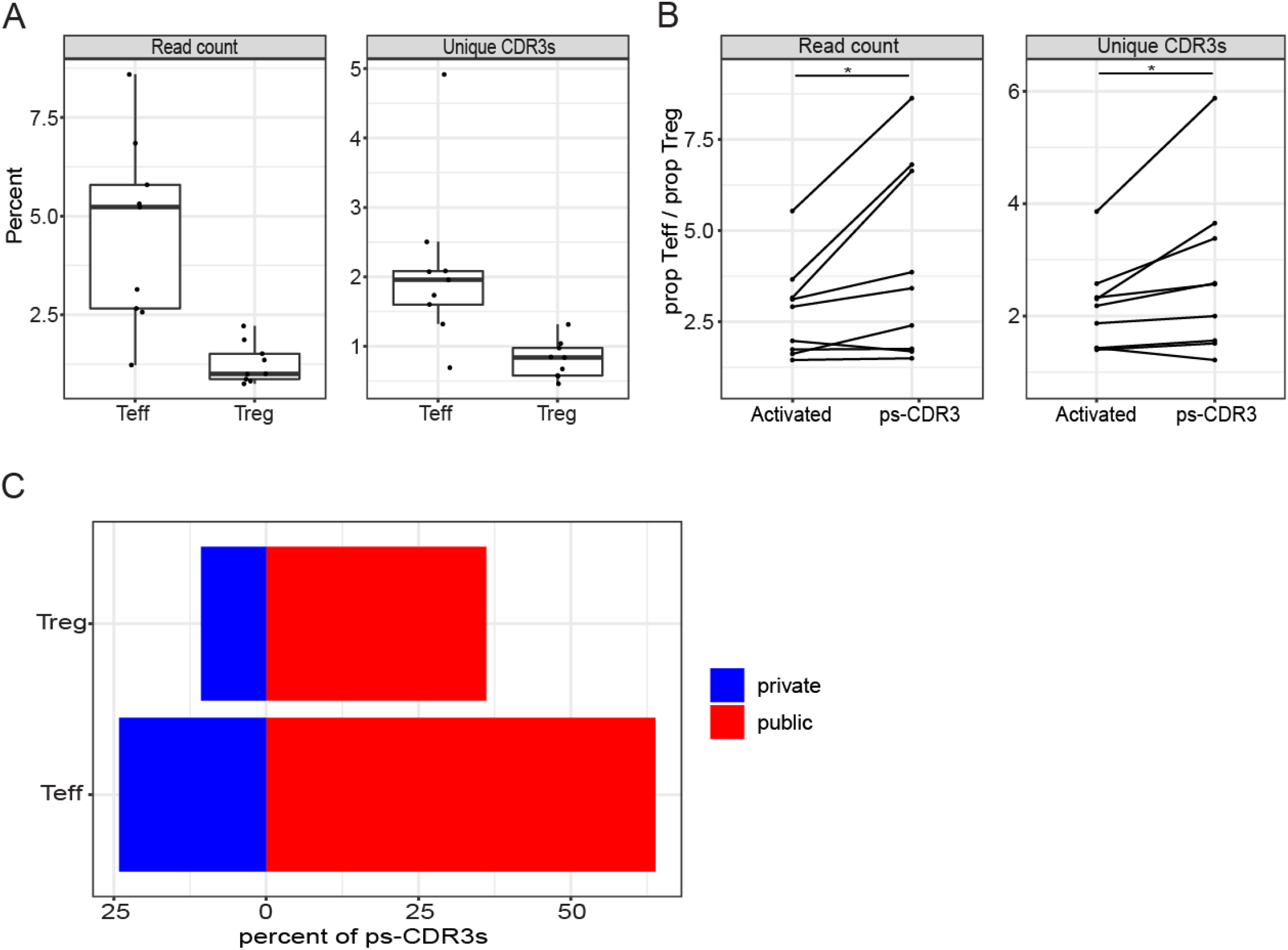
ps-CDR3s are more likely to be derived from T_eff_ than from T_reg_ in peanut-allergic individuals. A, The percentage of both unique CDR3s and total read counts corresponding to a ps-CDR3 sequence was higher in T_eff_ samples than in T_reg_ samples (n = 9). B, The disbalance between T_eff_-derived ps-CDR3s and T_reg_-derived ps-CDR3s was enhanced by our statistical enrichment. Plots show the proportion of T_eff_ ps-CDR3s : T_reg_ ps-CDR3s when using either the entire CD154^+^ CDR3 pool or the ps-CDR3s as probes. Proportions corresponding to both total reads (left) and unique sequences (right) are shown (*P < 0.05, paired Wilcoxon signed-rank test). C, Public ps-CDR3s were more likely to be found in the T_eff_ and T_reg_ subset repertoires than private ps-CDR3s. The percentage of public ps-CDR3s detected in any of the T_eff_ (bottom) or T_reg_ (top) samples was significantly higher than that of the private ps-CDR3s (P < 0.01, Fisher’s exact test).

## Discussion

Selection of antigen-specific T cells based on their ability to bind an MHC-peptide complex (pMHC-multimer selection) is a widely used technique, but this method depends on a priori knowledge of T cell epitopes in subjects with specific HLA backgrounds, and is therefore limited in scope. Here, we expand upon a method not constrained by these limitations, utilizing a biological enrichment and augment it with a statistical enrichment to isolate likely antigen-specific CDR3s that are not derived from bystander-activated T cells. This approach generated a library of ps-CDR3s that exhibited properties of an antigen-specific pool in terms of diversity, homology and convergence. Using these features in a network analysis to cluster similar sequences, we were able to show that public ps-CDR3s tended to have more neighbors (i.e. were more similar) than private ps-CDR3s, which corresponds with observations made by Madi et al. [22]. When using the ps-CDR3s to probe the T cell subset repertoires of 9 peanut-allergic individuals, we detected more T_eff_ ps-CDR3s than T_reg_ ps-CDR3s, indicating that an overrepresented peanut-specific T_eff_ repertoire is a characteristic of peanut allergy.

The use of CD154 as a marker to isolate antigen-activated T cells was first described in 2005 when Frentsch et al. demonstrated the utility of this method with an array of microbial and allergenic antigens [23]. Since then, numerous other groups have successfully applied this method with allergens and pathogen-derived proteins to better describe the phenotypes of antigen-specific CD4^+^ T cell populations [13] [24] [25] [26] [9] [27]. However, applying this methodology to better characterize antigen-specific TCR repertoires is in its infancy. Glanville et al. single-cell sorted and sequenced the TCRs of CD154^+^ T cells activated with an *M. tuberculosis* lysate and successfully validated antigen-specific motifs with an *in vitro* binding assay, demonstrating that CD154 up-regulation is a valid method to study antigen-specific repertoires [2]. Given the concern of bystander activation of non-antigen-specific T cells, we chose to use the CD154 up-regulation assay in conjunction with a statistical enrichment approach to define our library of ps-CDR3s. This statistical enrichment has the distinct advantage of utilizing autologous resting memory TCR repertoires as a reference when selecting the ps-CDR3s, hereby controlling for overrepresented clones in the CD154^+^ compartment due to bystander activation. The ps-CDR3s exhibited characteristics associated with antigen-specific TCR pools, such as increased homology and decreased diversity, when compared to total activated CDR3s as well as resting CDR3s. These findings emphasize the utility of applying the biological (CD154 up-regulation) and statistical (*G*-test of independence) enrichments in combination when determining the most relevant TCRs. Utilizing this approach in conjunction with single-cell technologies in future studies would allow for the analysis of paired αβ-TCR sequences and enhanced functional validations.

The application of this novel method for defining antigen-specific ps-CDR3s has the potential to answer a wide array of biological questions. As described here, antigen-specific ps-CDR3 libraries can be used to probe phenotypically defined subsets of CD4^+^ T cells to better understand the phenotype and physiology of antigen-specific T cells. Recently, our group has successfully used this approach to identify differences in the peanut-specific CD4^+^ T cell repertoire and phenotypes between peanut-allergic individuals with high and low clinical sensitivity [14]. We found that high clinical sensitivity correlates with an expanded and more diverse antigen-specific T_eff_ cell population, rather than a lack of T_reg_ cells. Our strategy could also be extended to track antigen-specific ps-CDR3s across a time-course or therapy. Examining the clones that expand or contract in response to an intervention could lead to insights regarding efficacy or susceptibility to side-effects.

A major advantage of our approach is that it is unbiased in terms of MHC-presented epitopes, representing the overall CD4^+^ T cell response to an antigen that is unlikely to be captured with pMHC-multimer-based sorting. In a previous study, Ryan et al. used a pMHC-dextramer approach to capture Ara h 2-specific T cells from peanut allergic individuals with a defined HLA-background (HLA-DRB1*1501, HLA-DRB4) [12]. While this method was fairly successful in identifying phenotypic changes in Ara h 2-specific T cells over the course of peanut oral immunotherapy, the authors were only able to define 13 Ara h 2-specific TCRs, providing limited insights into the peanut-specific repertoire. Using a method agnostic to both HLA-genotype and known epitopes can help better define the TCR landscape of a given immune response. For example, our study enabled us to confidently describe the public nature of 36 unique antigen-specific CDR3s. This would have been difficult to capture with methods based on pMHC-multimers, given the very low numbers of antigen-specific T cells detected by these methods and the sparsity of these T cells in the overall T cell repertoire. The propensity of the public ps-CDR3s to demonstrate convergent recombination further exemplifies the utility of our methodology, as convergence suggests antigenic selection [28] [7] [29].

In sum, by using a combination of biological and statistical enrichment of CDR3s from antigen-activated CD4^+^ T cells, we were able to identify private and public CDR3s that show evidence of antigenic selection. This method holds promise for application in a wide range of T cell-mediated disorders, as it effectively interrogates the antigen-specific TCR repertoire in unselected subjects, and potentially yields biomarkers for disease state and response to various treatments.

## Methods

### Subjects

The subjects included in this study participated in a peanut oral immunotherapy trial (NCT01750879) conducted at the Food Allergy Center at Massachusetts General Hospital. Criteria to be screened for the study included having a previous diagnosis of peanut allergy, a history of peanut-induced reactions consistent with immediate hypersensitivity and peanut-specific serum IgE levels greater than 5 kU/L (ImmunoCAP; Thermo Fisher). Subjects who met these criteria then underwent a double-blind placebo-controlled food challenge to confirm peanut allergy. Increasing peanut protein doses were administered every 20 minutes to a maximum dose of 300 mg according to the following schedule: 3, 10, 30, 100, and 300 mg. All 27 individuals included in this study had an objective allergic reaction during peanut challenge.

### Cell culture and sorting of peanut-activated and -resting memory CD4^+^ T cells

Patient blood samples were collected at baseline (before the start of peanut oral immunotherapy), and PBMCs were isolated by means of density gradient centrifugation (Ficoll-Paque Plus; GE Healthcare, Chicago, Ill). Fresh PBMCs were cultured in AIM V medium (Gibco, Waltham, Mass) for 20 hours at a density of 5 × 10^6^ in 1 mL medium per well in 24-well plates, and were left unstimulated or cultured with 100 μg/mL peanut protein extract (15 ×10^6^ PBMCs per variable). The peanut extract was prepared by agitating defatted peanut flour (Golden Peanut and Tree Nuts, Alpharetta, Ga) with PBS, centrifugation, and sterile-filtering.

Phycoerythrin-conjugated anti-CD154 (clone TRAP1; BD Biosciences, San Jose, Calif) was added to the cultures (20 μL/well) for the last 3 hours. After harvesting, the cells were labeled with AF700-conjugated anti-CD3 (clone UCHT1), allophycocyanin Cy7–conjugated anti-CD4 (RPA-T4), fluorescein isothiocyanate–conjugated anti-CD45RA (HI100), phycoerythrin-conjugated anti-CD154 (all from BD Biosciences), AF647-conjugated anti-CD69 (FN50; BioLegend, San Diego, Calif), and Live/Dead Fixable Violet stain (L34955; Thermo Fisher). Live CD3^+^CD4^+^CD45RA^−^ activated CD154^+^ and resting CD154^−^CD69^−^ T cells were sorted with a FACSAria II instrument (BD Biosciences). Sorted T cells were lysed in Buffer RLT Plus (Qiagen, Hilden, Germany) with 1% β-mercaptoethanol (Millipore-Sigma, St Louis, Mo), and stored at −80C, before total RNA and genomic DNA were isolated using the AllPrep DNA/RNA Micro Kit (Qiagen).

### TCRβ sequencing

Genomic DNA was used to amplify and sequence the complementarity-determining region 3 (CDR3) regions (immunoSEQ assay; Adaptive Biotechnologies, Seattle, Wash). The immunoSEQ approach generates an 87 basepair fragment capable of identifying the VDJ region spanning each unique CDR3. Amplicons were sequenced using the Illumina NextSeq platform. Using a baseline developed from a suite of synthetic templates, primer concentrations and computational corrections were used to correct for the primer bias common to multiplex PCR reactions. Raw sequence data were filtered on the basis of TCRβ V, D, and J gene definitions provided by the IMGT database (www.imgt.org) and binned using a modified nearest-neighbor algorithm to merge closely related sequences and remove both PCR and sequencing errors.

### Selection of ps-CDR3s

TCRβ sequencing data from activated (CD154^+^) and resting (CD154^−^CD69^−^) memory CD4^+^ T cell samples of all 27 peanut-allergic individuals was parsed using the *tcR* package (version 2.2.3) in R [30]. We applied a statistical method to define putatively peanut-specific CDR3s (ps-CDR3s), i.e. sequences that are likely to be antigen-specific and not detected due to bystander activation. This methodology’s workflow starts with performing a G-test of independence on every unique CDR3 found in an individual’s activated CD154^+^ T cell sample, to determine if the proportion of a given sequence is higher in the activated CDR3s versus the resting CDR3s. A G-test (also known as a likelihood ratio test) can be described by the formula:

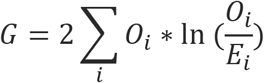

where *O_i_* is the observed read count of a given CDR3 in a particular population (activated or resting) and *E_i_* is the expected count based on the proportion of this CDR3 in the entire memory T cell population (pooled activated and resting CDR3s). With the G value for each CDR3, the probability that the proportion of a CDR3 in the activated compartment is derived by chance from the proportion in the resting compartment can be calculated. Given the sizable number of CDR3s that were analyzed from each individual’s data, a false discovery rate (FDR) correction was used to generate adjusted P-values (q-value) for each sequence, and those that met a cutoff of q < 0.05 were considered significant for this study. To enhance the stringency and reduce type 1 error, we also filtered out those CDR3s with a read count of only 1 in the activated compartment (n = 24520, 76.9%) and those with a proportion in the activated compartment being less than that in the resting compartment (n = 4, 0.01%). All sequences selected after filtering were deemed putatively peanut-specific (ps−)CDR3s.

### Minimum Hamming and Levenshtein distances

To determine global levels of similarity, the minimum Hamming distance (number of amino acid differences among CDR3s of same length) and minimum Levenshtein distance (minimum number of insertions/deletions/substitutions between CDR3s) of each ps-CDR3 against all other ps-CDR3s was determined using R (version 3.5.1) with the package *stringdist* (version 0.9.5.1). The percentage of ps-CDR3s at each minimum Hamming or Levenshtein distance was calculated. As a control, the minimum Hamming and Levenshtein distances of 100 equal-sized random resamplings of CDR3s from the total activated and resting T cell pools were determined. The median percentage of sequences at each minimum Hamming and Levenshtein distance for the 100 resamplings was calculated.

### Diversity measurements

Diversity curves that measured Hill’s diversity metric across diversity orders 0-4 were created using the R package *alakazam* (version 0.3.0) with the *alphaDiversity* function [17]. Hill’s diversity metric can be described by the following formula:

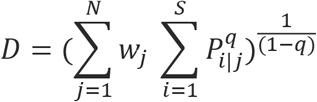

Where *i* represents a unique CDR3, *j* is the sample size, *P*_*i|j*_ is the proportional abundance of the *i*^*th*^ CDR3 in the *j*^*th*^ sample, *W*_*j*_ is the proportional abundance of the *j*^*th*^ sample relative to the entire dataset and *q* is a tuning parameter. *q* controls the influence of abundant over rare species on the metric (as *q* increases, the abundance of dominant sequences is more heavily weighted). Therefore, examining Hill’s diversity metric across different values of *q* captures both the richness and abundance of species. The fold-change of Hill’s diversity metric of all pairwise combinations of ps-CDR3s, total activated CDR3s and resting CDR3s was determined and the differences in these distributions were compared.

### Motif analysis

CDR3 amino acid sequences were trimmed to the region most likely to be in contact with antigenic peptide (IMGT positions 106-117), as the stem positions of CDR3s are not predicted to be involved with antigen binding [2]. The trimmed sequences were then broken into all possible continuous nmers of size 3, 4, and 5 amino acids. In addition, discontinuous motifs were generated of size 4 and 5, allowing for gaps in the sequences so long as there were still 3 conserved residues. The proportion of each nmer in the ps-CDR3s was calculated by dividing the number of reads containing that nmer by the total number of ps-CDR3 reads. To determine fold-enrichment, this proportion of ps-CDR3 reads with a given nmer was divided by the proportion of reads in the resting CDR3s with the nmer.

### Network analysis

Each unique ps-CDR3 sequence was represented as a node and edges between nodes were made if there was a Levenshtein distance of 1 between them or if they shared an enriched motif. Levenshtein distances were determined using the R package *stringdist* (version 0.9.5.1). A self-edge was created on a node for every additional nucleotide sequence that corresponded to the same amino acid sequence. Network object (gml) files were created in R using the *igraph* package (version 1.0.1) and network visualization was performed with Cytoscape 3.7.0, using a force-directed open-CL layout. To evaluate the amount of structure present among ps-CDR3s, the number of edges in ps-CDR3s was compared with the median number of edges in 100 equal-sized random resamplings of resting and activated CDR3 sequences, by creating edges between the sequences of each resampling if they were within a Levenshtein distance of 1, had an enriched motif or if sequences demonstrated convergence.

### T cell subset analysis

Cryopreserved PBMCs from 9 of the 27 subjects that were used to generate the ps-CDR3 library were thawed and labeled with BV650-conjugated anti-CD3 (UCHT1), phycoerythrin-Cy7-conjugated anti-CD4 (RPA-T4), allophycocyanin-H7–conjugated anti-CD45RA (HI100) (all from BD Biosciences), BV605-conjugated anti-CD25 (BC96), BV785-conjugated anti-CD127 (A019D5) (both from BioLegend), and Live/Dead Fixable Blue stain (L23105; Thermo Fisher). Live CD3^+^CD4^+^CD45RA^−^ CD25^+^CD127^+^ T_eff_ and CD25^++^CD127^−^ T_reg_ cells were sorted with a FACSAria Fusion instrument (BD Biosciences). Sorted T cell subsets then underwent the TCRβ sequencing protocol described above. These T_eff_ and T_reg_ libraries were probed for amino acid sequences also found in the ps-CDR3 pool.

### Statistics

All comparisons made with matched pairs of values from individual subjects were tested with a Wilcoxon matched-pairs signed rank test. Such comparisons include frequencies of CD154^+^ T cells per million CD4^+^ T cells in cultures, differences in the ratios of diversities in resting, activated and ps-CDR3 pools, as well as dysbalance of T_eff_:T_reg_ with and without statistical filtering. Evaluation of enrichment of ps-CDR3s at minimum Hamming/Levenshtein distances utilized a 2-sided Fisher’s exact test, comparing the median frequencies of activated or resting CDR3s to the observed frequency in the ps-CDR3s. To assess differences in diversity, the empirical cumulative distribution function of bootstrap delta distribution was used as described by Stern et al. [31]. Comparisons of public and private ps-CDR3 repertoire features, such as the number of nucleotide insertions, total rearrangements and node degree in the ps-CDR3 network were done with a Mann-Whitney U test. A 2-sided Fisher’s exact test was used to compare the proportion of public ps-CDR3s with an enriched motif to private ps-CDR3s with a motif. Similarly, a 2-sided Fisher’s exact test was used to compare the proportion of public ps-CDR3s with an edge in the ps-CDR3 network to that of the private ps-CDR3s. A z-test was used to evaluate the number of edges created within the ps-CDR3 network to that of the distribution of edges created by 100 equal-sized random samplings of resting or activated CDR3s. A two-sided Fisher’s exact test was used to compare the proportion of public ps-CDR3s found in the T cell subsets to private ps-CDR3s in T cell subsets.

### Study approval

All subjects were recruited with informed consent prior to sample collection, and the study was approved by the Institutional Review Board of Partners Healthcare (protocol no. 2012P002153). The study was conducted according to the principles of the Declaration of Helsinki.

## Author contributions

NPS, BR, and WGS designed the study. NPS and BR performed all of the lab work and generated all of the data. NPS, YVV and WGS performed the data analysis. NPS and BR wrote the manuscript. All authors reviewed and approved the final version of the manuscript.

## Acknowledgements

We would like to thank Lauren Tracy, Colby Rondeau, Christine Elliot, and Leah Hayden, the clinical research coordinators who were instrumental in obtaining the samples used in the study. We would also like to thank the staff at the MGH Department of Pathology Flow and Image Cytometry Research Core who helped with the FACS. The Flow Core obtained funding from the National Institutes of Health Shared Instrumentation program (grant nos. 1S10OD012027-01A1, 1S10OD016372-01, 1S10RR020936-01, and 1S10RR023440-01A1). This work was supported by the NIH/NIAID (grant U19AI095261 to WGS. and Andrew D. Luster, and K23AI130408 to YVV) and the Food Allergy Science Initiative (FASI).

---

**Supplemental figure 1.**
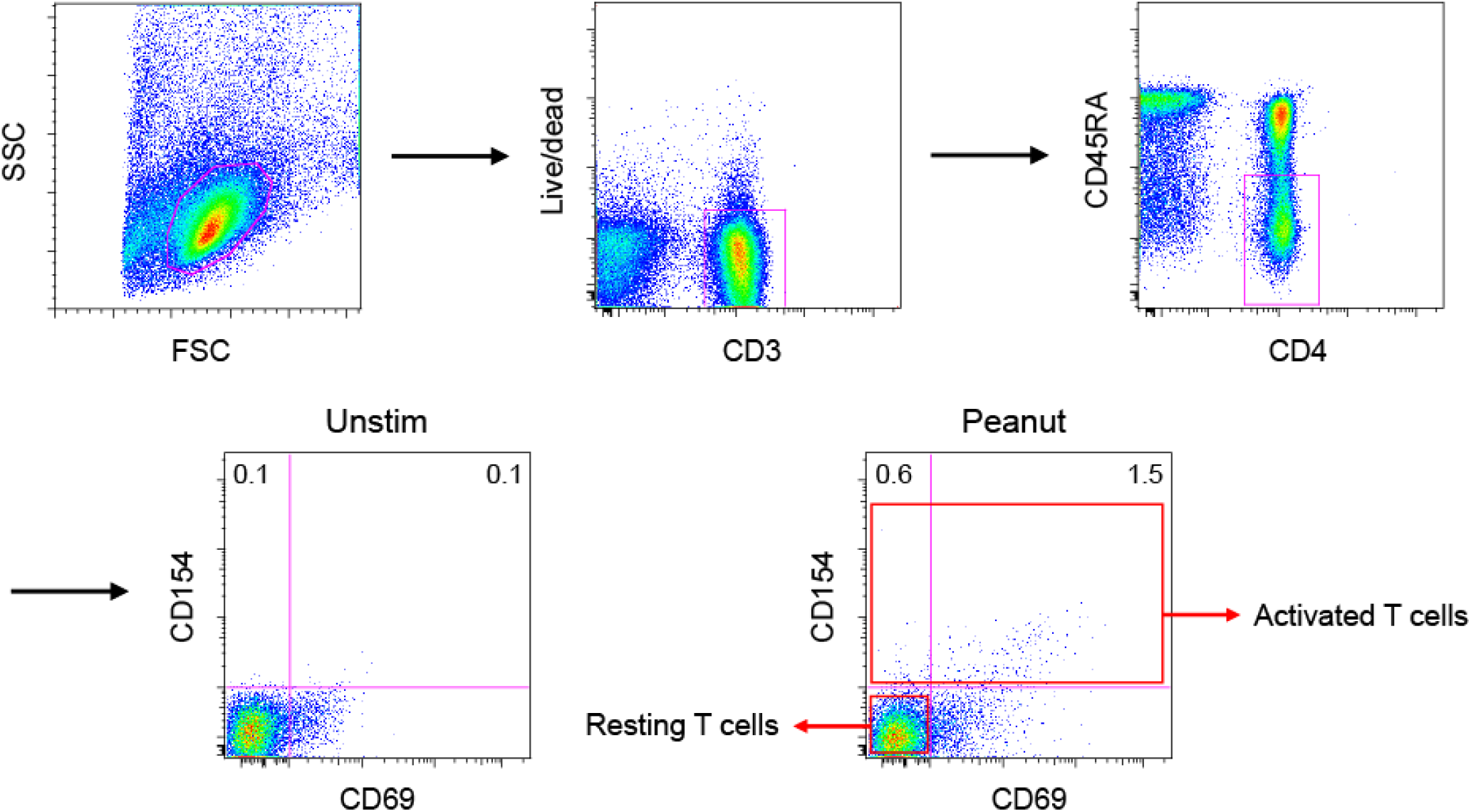
FACS gating strategy for isolating peanut-activated memory CD4^+^ T cells (CD154^+^) and resting memory CD4^+^ T cells (CD154^−^CD69^−^) from the PBMCs of peanut-allergic individuals.

**Supplemental figure 2.**
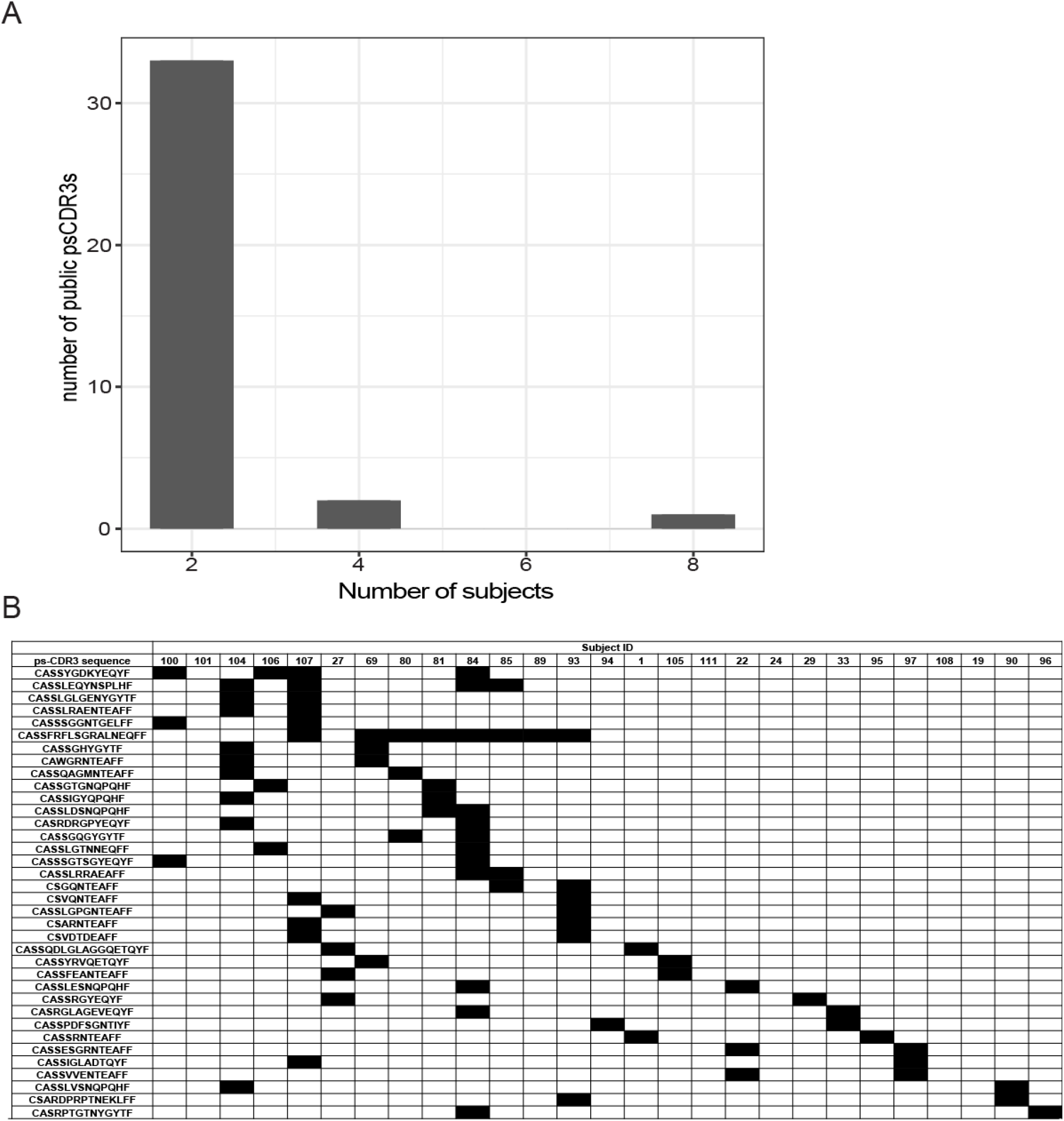
Distribution of public ps-CDR3s across peanut-allergic subjects. A, Distribution of the public ps-CDR3s over the patients. Shown is the number of public ps-CDR3s present in a given number of patients. B, Summary of the presence of each unique public ps-CDR3 in each subject.

**Supplemental figure 3.**
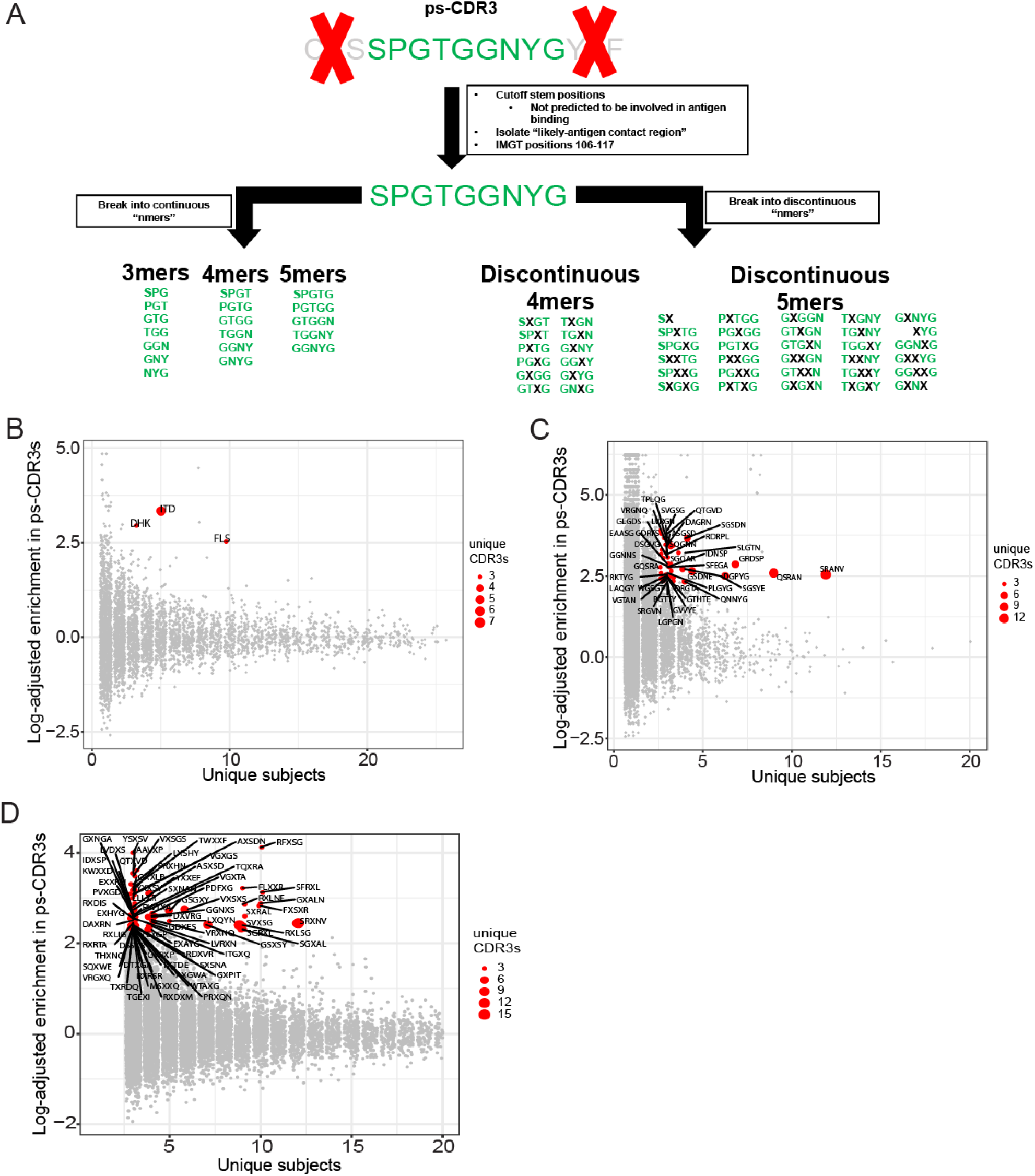
Motif analysis of ps-CDR3s. A, Schema of nmer generation with an example ps-CDR3 (see methods). B-D, The ps-CDR3s contain shared and enriched 3-mer, 5- mer, and discontinuous 5-mer motifs as compared to resting CDR3s. Red points represent the most dominant motifs, which were ≥10-fold enriched, in ≥ 3 unique ps-CDR3s derived from ≥ 3 individuals.

**Supplemental figure 4.**
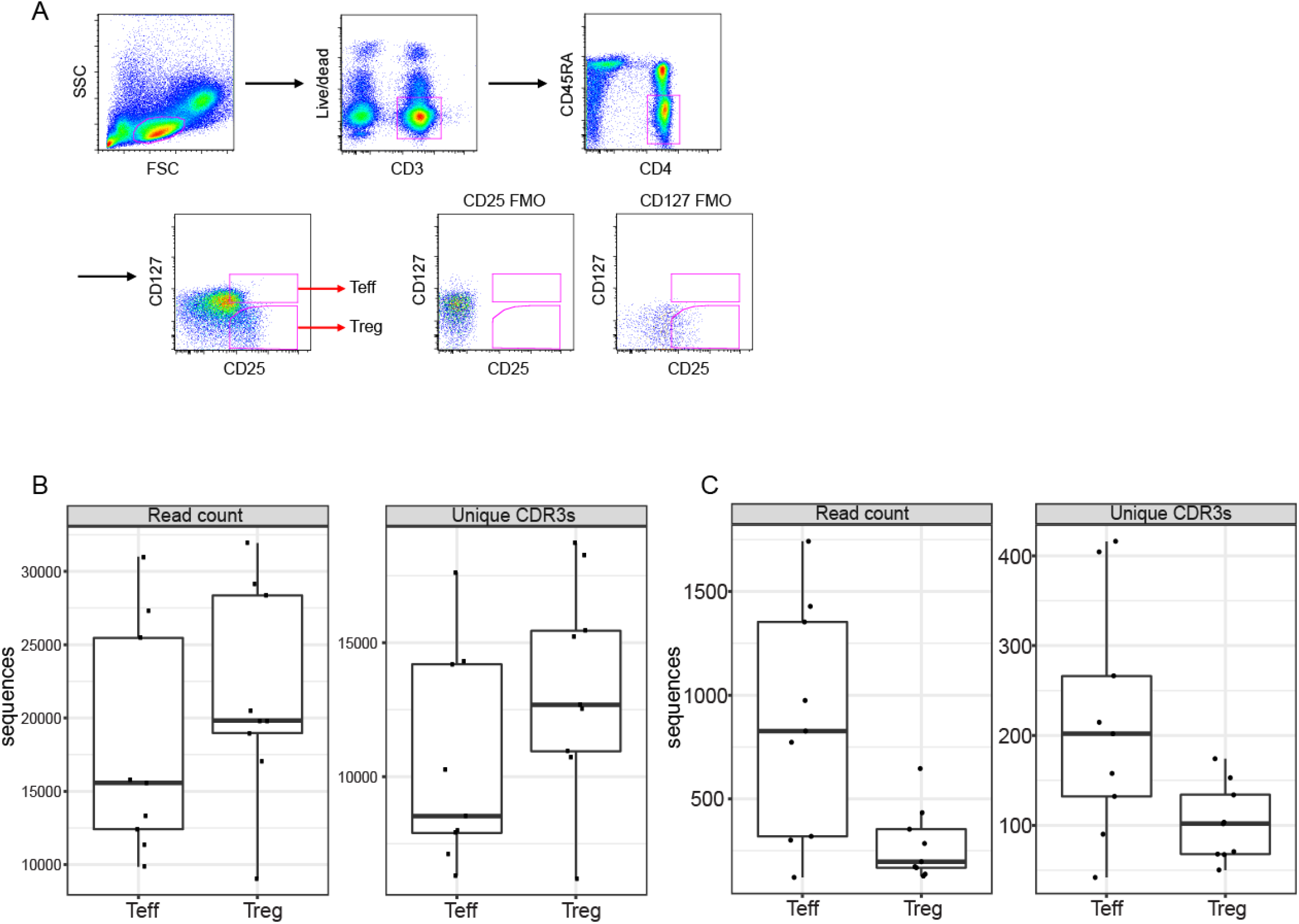
Bulk-isolation of T_eff_ and T_reg_, and counts of total CDR3s. A, FACS gating strategy for isolating bulk memory CD4^+^ T_eff_ (CD25^+^CD127^+^) and memory CD4^+^ T_reg_ (CD25^++^ CD127^−^) cells. B, Overall, reads and unique CDR3s were captured from T_reg_ samples than T_eff_ samples (n = 9). C, The absolute number of both unique and total read counts of ps-CDR3s was higher in T_eff_ than in T_reg_ (n = 9).

**Supplemental Table 1:** HLA genotypes of subjects in the study.

| Subject ID | DPA1_allele1 | DPA1_allele2 | DPB1_allele1 | DPB1_allele2 | DQA1_allele1 | DQA1_allele2 | DQB1_allele1 | DQB1_allele2 | DRB1_allele1 | DRB1_allele2 | DRB3_allele1 | DRB3_allele2 | DRB4 | DRB5 |
| --- | --- | --- | --- | --- | --- | --- | --- | --- | --- | --- | --- | --- | --- | --- |
| 1 | 1:03:01 | 2:02:02 | 1:01:01 | 3:01:01 | 1:03:01 | 3:03:01 AM | 3:01:01 | 6:03:01 | 13:01:01 | 4:01:01 | 1:01:02 |  | 1:03:01 |  |
| 19 | 2:01:01 | 2:07:01 | 11:01:01 | 19:01:01 | 1:03:01 | 2:01:01 | 2:02:01 | 6:03:01 | 13:01:01 | 7:01:01 | 2:02:01 |  | 1:01:01 |  |
| 22 | 1:03:01 | 1:03:01 | 4:01:01 | 4:01:01 | 1:02:01 | 3:01:01 | 3:02:01 | 6:02:01 | 15:01:01 | 4:02:01 |  |  | 1:03:01 | 1:01:01 |
| 24 | 2:01:01 | 2:01:02 | 1:01:01 | 17:01:01 | 2:01:01 | 5:01:01 | 2:01:01 | 2:02:01 | 3:01:01 | 7:01:01 | 1:01:02 |  | 1:01:01 |  |
| 27 | 1:03:01 | 2:01:01 | 3:01:01 | 10:01:01 | 1:04:01 | 3:01:01 | 3:02:01 | 5:03:01 | 14:54:01 | 4:01:01 | 2:02:01 |  | 1:03:01 |  |
| 29 | 1:03:01 | 2:01:01 | 4:01:01 | 13:FNVU | 3:01:01 | 5:05:01 | 3:01:01 | 3:02:01 | 11:04:01 | 4:02:01 | 2:02:01 |  | 1:03:01 |  |
| 33 | 1:03:01 | 2:01:07 | 4:01:01 | 13:FNVU | 1:02:01 | 3:02:01 | 3:03:02 | 6:09:01 | 13:02:01 | 09:CWA | 3:01:01 |  | 1:03:02 |  |
| 69 | 1:03:01 | 1:03:01 | 2:01:02 | 13:FNVU | 1:02:01 | 2:01:01 | 2:02:01 | 6:02:01 | 15:01:01 | 7:01:01 |  |  | 1:01:01 | 1:01:01 |
| 80 | 1:03:01 | 2 | 4:01:01 | 85:01:01 | 2:01:01 | 5:01:01 | 2:01:01 | 2:02:01 | 3:01:01 | 7:01:01 | 1:01:01 |  | 1:03:01 |  |
| 81 | 1:03:01 | 1:03:01 | 2:01:01 | 4:01:01 | 1:02:01 | 3:01:01 | 3:02:01 | 6:02:01 | 15:01:01 | 4:02:01 |  |  | 1:03:01 | 1:01:01 |
| 84 | 1:03:01 | 2:01:01 | 4:01:01 | 10:01:01 | 1:01:02 | 1:02:01 | 5:01:01 | 6:09:01 | 1:02:01 | 13:02:01 | 3:01:01 | 3:01:01 |  |  |
| 85 | 1:03:01 | 1:03:01 | 4:01:01 | 4:02:01 | 5:01:01 | 5:01:01 | 2:01:01 | 2:01:01 | 3:01:01 | 3:01:01 | 1:01:02 | 1:01:02 |  |  |
| 89 | 1:03:01 | 2:06 | 4:02:01 | 5:01:01 | 1:02:01 | 1:02:01 | 6:02:01 | 6:02:01 | 15:01:01 | 15:01:01 |  |  |  | 1:01:01 |
| 90 | 1:03:01 | 1:03:01 | 4:01:01 | 4:01:01 | 5:01:01 | 5:05:01 | 2:01:01 | 3:01:01 | 3:01 | 11:01 | 1:01:02 | 2:02:01 |  |  |
| 93 |  |  | 4:01:01 | 4:01:01 | 1:04:01 | 5:01:01 | 2:01:01 | 5:03:01 | 3:01:01 | 14:01:01 | 1:01:02 | 2:24 |  |  |
| 94 | 1:03:01 | 1:03:01 | 2:01:02 | 4:01:01 | 1:01:01 | 1:02:01 | 5:01:01 | 6:02:01 | 1:03:01 | 15:01:01 |  |  |  | 1:01:01 |
| 95 | 1:03:01 | 1:03:01 | 4:01:01 | 4:01:01 | 1:02:01 | 5:05:01 | 3:01:01 | 6:02:01 | 15:01:01 | 11:01:01 | 2:02:01 |  |  | 1:01:01 |
| 96 | 1:03:01 | 2:01:02 | 1:01:01 | 2:01:02 | 3:03:01 | 5:05:01 | 3:01:01 | 3:02:01 | 12:JV | 4:01:01 | 2:02:01 |  | 1:03:01 |  |
| 97 | 1:03:01 | 1:03:01 | 2:01:02 | 48:01:00 | 3:01:01 | 5:03:01 | 3:01:01 | 3:02:01 | 13:12:01 | 4:03:01 | 2:02:01 |  | 1:03:01 |  |
| 100 | 1:03:01 | 1:03:01 | 2:01:02 | 4:01:01 | 5:05:01 | 5:05:01 | 3:01:01 | 3:01:01 | 11:04:01 | 11:04:01 | 2:02:01 | 2:02:01 |  |  |
| 101 | 1:03:01 | 1:03:01 | 3:01:01 | 4:01:01 | 1:02:01 | 1:02:01 | 6:02:01 | 6:04:01 | 15:01:01 | 13:02:01 | 3:01:01 |  |  | 1:01:01 |
| 104 | 1:03:01 | 1:03:01 | 4:01:01 | 4:01:01 | 1:02:01 | 2:01:01 | 3:03:02 | 6:02:01 | 15:01:01 | 7:01:01 |  |  | 01:03:01N | 1:01:01 |
| 105 | 1:03:01 | 2:01:01 | 2:01:02 | 5:01:01 | 3:01:01 | 5:01:01 | 2:01:01 | 3:02:01 | 3:01:01 | 4:02:01 | 1:01:02 |  | 1:03:01 |  |
| 106 | 1:03:01 | 1:03:01 | 4:01:01 | 104:01:01 | 1:01:02 | 2:01:01 | 3:01:01 | 5:01:01 | 1:02:01 | 7:01:01 |  |  | 1:01:01 |  |
| 107 | 1:03:01 | 1:03:01 | 4:01:01 | 20:01:01 | 3:01:01 | 5:01:01 | 2:01:01 | 3:02:01 | 3:01:01 | 4:04:01 | 1:01:02 |  | 1:03:01 |  |
| 108 | 1:03:01 | 1:03:01 | 3:01:01 | 4:02:01 | 1:02:01 | 5:01:01 | 2:01:01 | 6:04:01 | 3:01:01 | 13:02:01 | 1:01:02 | 3:01:01 |  |  |
| 111 | 1:03:01 | 1:03:01 | 2:01:02 | 4:01:01 | 1:03:01 | 5:05:01 | 3:01:01 | 6:03:01 | 13:01:01 | 13:05:01 | 1:01:02 | 2:02:01 |  |  |

## References

1. Lythe, G., et al., How many TCR clonotypes does a body maintain? J Theor Biol, 2016. 389: p. 214–24.

2. Glanville, J., et al., Identifying specificity groups in the T cell receptor repertoire. Nature, 2017. 547(7661): p. 94–98.

3. Shugay, M., et al., VDJdb: a curated database of T-cell receptor sequences with known antigen specificity. Nucleic Acids Res, 2018. 46(D1): p. D419–D427.

4. Dash, P., et al., Quantifiable predictive features define epitope-specific T cell receptor repertoires. Nature, 2017. 547(7661): p. 89–93.

5. Burrows, S.R., et al., T cell receptor repertoire for a viral epitope in humans is diversified by tolerance to a background major histocompatibility complex antigen. J Exp Med, 1995. 182(6): p. 1703–15.

6. Li, H., et al., Determinants of public T cell responses. Cell Res, 2012. 22(1): p. 33–42.

7. Venturi, V., et al., Sharing of T cell receptors in antigen-specific responses is driven by convergent recombination. Proc Natl Acad Sci U S A, 2006. 103(49): p. 18691–6.

8. Sicherer, S.H. and H.A. Sampson, Food allergy: A review and update on epidemiology, pathogenesis, diagnosis, prevention, and management. J Allergy Clin Immunol, 2018. 141(1): p. 41–58.

9. Wambre, E., et al., A phenotypically and functionally distinct human TH2 cell subpopulation is associated with allergic disorders. Sci Transl Med, 2017. 9(401).

10. Gowthaman, U., et al., Identification of a T follicular helper cell subset that drives anaphylactic IgE. Science, 2019. 365(6456).

11. Syed, A., et al., Peanut oral immunotherapy results in increased antigen-induced regulatory T-cell function and hypomethylation of forkhead box protein 3 (FOXP3). J Allergy Clin Immunol, 2014. 133(2): p. 500–10.

12. Ryan, J.F., et al., Successful immunotherapy induces previously unidentified allergen-specific CD4+ T-cell subsets. Proc Natl Acad Sci U S A, 2016. 113(9): p. E1286–95.

13. Chattopadhyay, P.K., J. Yu, and M. Roederer, Live-cell assay to detect antigen-specific CD4+ T-cell responses by CD154 expression. Nat Protoc, 2006. 1(1): p. 1–6.

14. Ruiter, B., et al., Expansion of the CD4(+) effector T-cell repertoire characterizes peanut-allergic patients with heightened clinical sensitivity. J Allergy Clin Immunol, 2020. 145(1): p. 270–282.

15. Renand, A., et al., Heterogeneity of Ara h Component-Specific CD4 T Cell Responses in Peanut-Allergic Subjects. Front Immunol, 2018. 9: p. 1408.

16. Akdis, M., et al., Immune responses in healthy and allergic individuals are characterized by a fine balance between allergen-specific T regulatory 1 and T helper 2 cells. J Exp Med, 2004. 199(11): p. 1567–75.

17. Gupta, N.T., et al., Change-O: a toolkit for analyzing large-scale B cell immunoglobulin repertoire sequencing data. Bioinformatics, 2015. 31(20): p. 3356–8.

18. Khosravi-Maharlooei, M., et al., Crossreactive public TCR sequences undergo positive selection in the human thymic repertoire. J Clin Invest, 2019. 129(6): p. 2446–2462.

19. Madi, A., et al., T-cell receptor repertoires share a restricted set of public and abundant CDR3 sequences that are associated with self-related immunity. Genome Res, 2014. 24(10): p. 1603–12.

20. Venturi, V., et al., TCR beta-chain sharing in human CD8+ T cell responses to cytomegalovirus and EBV. J Immunol, 2008. 181(11): p. 7853–62.

21. Ritvo, P.G., et al., High-resolution repertoire analysis reveals a major bystander activation of Tfh and Tfr cells. Proc Natl Acad Sci U S A, 2018. 115(38): p. 9604–9609.

22. Madi, A., et al., T cell receptor repertoires of mice and humans are clustered in similarity networks around conserved public CDR3 sequences. Elife, 2017. 6.

23. Frentsch, M., et al., Direct access to CD4+ T cells specific for defined antigens according to CD154 expression. Nat Med, 2005. 11(10): p. 1118–24.

24. Archila, L.D., et al., Ana o 1 and Ana o 2 cashew allergens share cross-reactive CD4(+) T cell epitopes with other tree nuts. Clin Exp Allergy, 2016. 46(6): p. 871–83.

25. Chiang, D., et al., Single-cell profiling of peanut-responsive T cells in patients with peanut allergy reveals heterogeneous effector TH2 subsets. J Allergy Clin Immunol, 2018. 141(6): p. 2107–2120.

26. Weissler, K.A., et al., Identification and analysis of peanut-specific effector T and regulatory T cells in children allergic and tolerant to peanut. J Allergy Clin Immunol, 2018. 141(5): p. 1699–1710 e7.

27. Stonier, S.W., et al., Marburg virus survivor immune responses are Th1 skewed with limited neutralizing antibody responses. J Exp Med, 2017. 214(9): p. 2563–2572.

28. Joachims, M.L., et al., Single-cell analysis of glandular T cell receptors in Sjogren’s syndrome. JCI Insight, 2016. 1(8).

29. Ruggiero, E., et al., High-resolution analysis of the human T-cell receptor repertoire. Nat Commun, 2015. 6: p. 8081.

30. Nazarov, V.I., et al., tcR: an R package for T cell receptor repertoire advanced data analysis. BMC Bioinformatics, 2015. 16: p. 175.

31. Stern, J.N., et al., B cells populating the multiple sclerosis brain mature in the draining cervical lymph nodes. Sci Transl Med, 2014. 6(248): p. 248ra107.

